# EEG biomarkers of reduced inhibition in human cortical microcircuits in depression

**DOI:** 10.1101/2021.07.18.452836

**Authors:** Frank Mazza, Taufik A. Valiante, John D. Griffiths, Etay Hay

## Abstract

Reduced cortical inhibition by somatostatin-expressing (SST) interneurons has been strongly associated with treatment-resistant depression. However, whether the effects of reduced SST interneuron inhibition on microcircuit activity have signatures detectible in electroencephalography (EEG) signals remains unknown. We simulated resting-state activity and EEG using detailed models of human cortical microcircuits with normal (healthy) or reduced SST interneuron inhibition (depression). Healthy microcircuit models showed emergent key features of resting-state EEG, and depression microcircuits exhibited increased theta, alpha and low beta power (4 – 15 Hz). The changes in depression involved a combination of an aperiodic broadband, and periodic theta components. We then demonstrated the specificity of the EEG signatures of reduced SST interneuron inhibition by showing they were distinct from those corresponding to reduced parvalbumin-expressing (PV) interneuron inhibition. Our study thus links SST interneuron inhibition level to distinct features in EEG simulated from detailed human microcircuits, which can serve to better identify mechanistic subtypes of depression using EEG, and non-invasively monitor modulation of cortical inhibition.

## Introduction

Major depressive disorder (depression) is a leading cause of disability worldwide[1] and involves varying mechanisms and symptoms[2,3]. Consequently, a significant proportion of patients remain resistant to antidepressants[4] and second-line treatments[5]. Electro-encephalography (EEG) offers a non-invasive and cost-effective method for brain signal-based biomarkers to improve diagnosis, monitoring and treatment of depression subtypes[6,7]. However, the multi-scale mechanisms of depression and their link to clinically-relevant brain signals remain poorly understood.

There is growing evidence that reduced cortical inhibition plays a mechanistic role in depression[8] and treatment-resistant depression[9–11], especially inhibition mediated by somatostatin-expressing (SST) interneurons[11–15]. Recent studies showed a marked reduction in SST expression by SST interneurons in post-mortem tissue of depression patients, across all layers of the prefrontal cortex (PFC) and anterior cingulate cortex, indicating a decrease in SST interneuron inhibition[16–18]. In rodent models, silencing SST interneuron inhibition led to depression symptoms[15], and novel therapeutic compounds acting via positive allosteric modulation of alpha-5-gamma-aminobutyric-acid-A (*α*5-GABA_A_) receptors targeted by SST interneurons resulted in pro-cognitive and anxiolytic effects[15,19]. In contrast, indications of changes in parvalbumin-expressing (PV) interneuron inhibition were inconsistent and less pronounced, signifying a more selective vulnerability for SST interneurons in depression[20,21]. Furthermore, other disorders such as Alzheimer’s and aging that show changes in SST interneuron inhibition involve other key changes such as cell and synapse loss, therefore the altered inhibition may play a less central or less consistent role in these conditions than in depression[22–24]. Changes in SST interneuron inhibition on circuit activity could have signatures detectible in EEG, due to this cell type’s principal role in modulating input to pyramidal (Pyr) neurons, which are major contributors to EEG signals along with L5/6[8,21]. SST interneurons provide synaptic and tonic inhibition onto the apical dendrites of Pyr neurons[27,28], and mediate lateral inhibition in the cortex[25,29,30], particularly during periods of quiet, resting wakefulness (resting state)[31]. Accordingly, previous studies indicate that reduced SST interneuron inhibition in depression increases baseline activity of Pyr neurons[12,25,32].

While the contribution of SST interneurons to resting-state cortical oscillations remains largely unknown, studies show a role for this cell type in modulating low-frequency oscillations. SST stimulation entrains network activity in the 5 – 30 Hz range, and SST suppression modulates theta band (4 – 8 Hz) power[33,34]. SST interneurons have also been suggested to govern network synchrony in slow-wave sleep, which is marked by slow oscillations[35]. Thus, reduced SST interneuron inhibition in depression may affect EEG low-frequency power. Conversely, PV interneurons have been shown to modulate high beta (20 – 30 Hz) and gamma (30 – 50 Hz) frequencies in human hippocampal LFP recordings, indicating that these two interneuron types likely modulate distinct frequency domains in recorded EEG[36,37]. We recently showed that a 40% reduction in SST interneuron inhibition, estimated from post-mortem tissue studies of depression patients[16], significantly increased baseline Pyr spike rate in simulated human microcircuits[32]. However, it is still unknown if this level of reduction in SST interneuron inhibition would significantly alter baseline oscillatory dynamics detectable in EEG.

Previous studies have linked neuronal spiking to extracellular signals, although mostly using animal models and local field potential (LFP)[38–40]. Human studies have characterized LFP oscillations in cortical slices and showed phase-amplitude coupling between deep and superficial layers[41]. Others have studied the phase preference of Pyr neuron spikes relative to spindle events in rodent intracranial EEG oscillations[42]. However, experimental methods are limited in their ability to characterize the effects of the cellular changes in depression on human brain signals *in vivo,* thus meriting the use of computational models. Previous computational studies have identified inter-laminar mechanisms underlying evoked related potentials during stimulus response using simplified neuron morphologies and connectivity[43,44]. The increased availability of human neuronal electrical and synaptic connectivity data [30,45,46] provides important constraints for detailed models of human cortical microcircuits[32], which can be used to link mechanisms of microcircuit activity in human health and disease to signatures in local circuit-generated EEG signals[47,48].

In this study, we identified EEG biomarkers of microcircuit effects due to reduced SST interneuron inhibition, as estimated from gene expression changes in depression. Using biophysically detailed models of human cortical microcircuits, we simulated resting-state activity in health and depression together with local EEG signals. We characterized changes in restingstate EEG spectral power, and in relation to spiking activity in different neuron types, to identify biomarkers of reduced SST interneuron inhibition in depression.

## Results

### Human cortical microcircuit models reproduce resting-state EEG features

We used our previous detailed models of human cortical L2/3 microcircuits[32] (Fig 1a) as canonical cortical microcircuit models for simulating resting state spiking and EEG signals. The model microcircuits included the four key neuron types: Pyr neurons, SST interneurons, Parvalbumin-expressing interneurons (PV), and Vasoactive intestinal polypeptide-expressing interneurons (VIP). To simulate intrinsic healthy resting-state spiking activity, all neurons received random background excitatory input corresponding to baseline cortical and thalamic drive. The model microcircuits were implemented in a physical volume, enabling simulation of LFP and EEG together with microcircuit spiking (Fig 1a-d).

**Figure 1.**
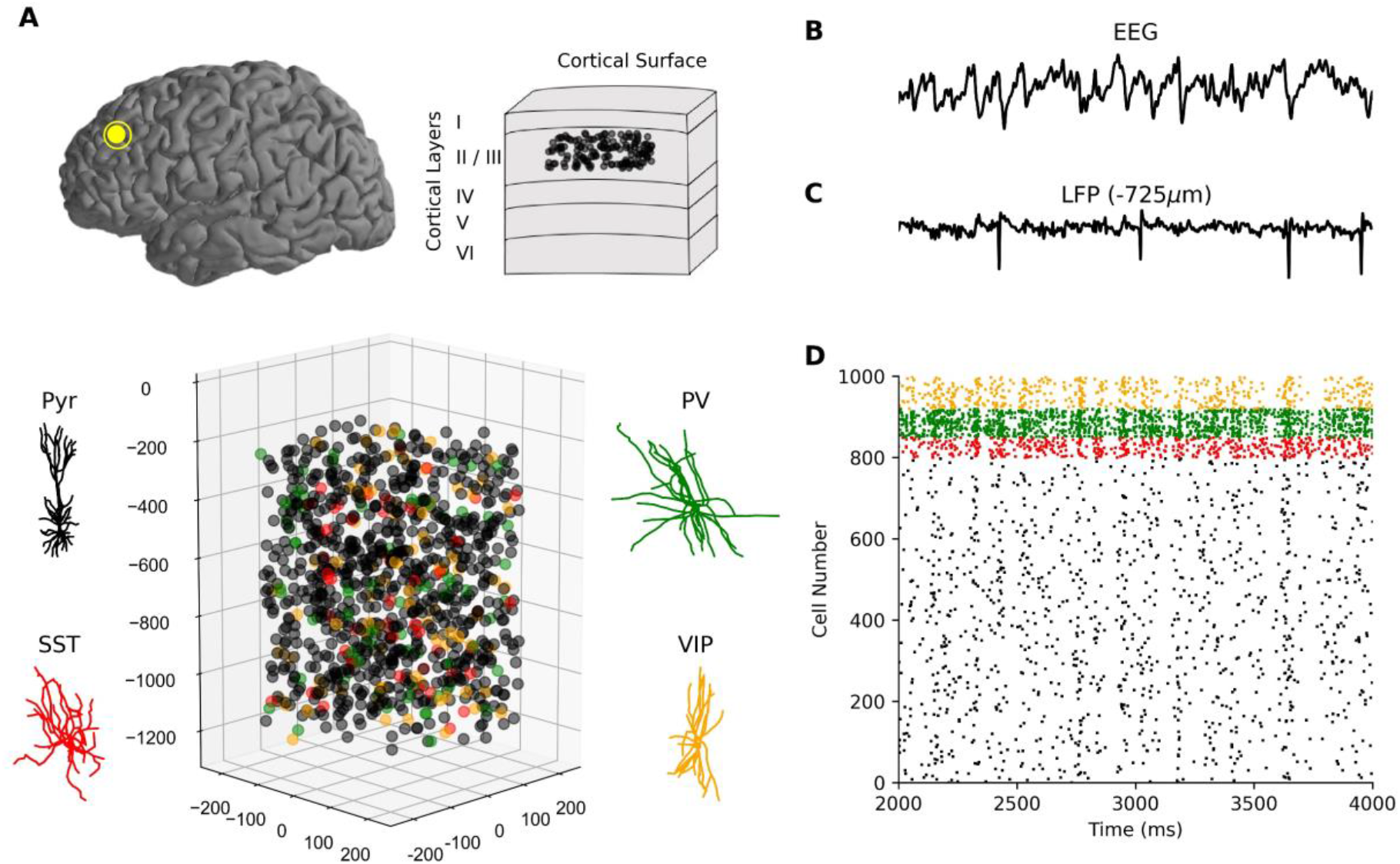
Simulating neuronal spiking and EEG signals from human cortical microcircuits. **(a)** Detailed models of human cortical microcircuits, showing somatic placement of 1000 neurons in a 500×500×950 μm^3^ volume along layer 2/3 (250 – 1200 μm below pia) and reconstructed morphologies used in the neuron models. **(b – d)** Temporally aligned multi-scale simulated signals: EEG (b) from scalp electrode directly above the microcircuit; LFP signal (c) recorded at the middle of L2/3 (depth of −725 μm); Raster plot of spiking in different neurons in the microcircuit (d), color-coded according to neuron type. Neurons received background excitatory inputs to generate intrinsic circuit activity.

Baseline microcircuit activity was oscillatory, marked by synchronous spiking events (Fig 1d) and corresponding fluctuations in LFP and EEG signals (Fig 1b, c). We quantified oscillatory activity in the healthy microcircuits by calculating EEG power spectral density (PSD). The microcircuit EEG exhibited a peak in theta (4 – 8 Hz) and alpha (8 – 12 Hz) bands (n = 60 randomized microcircuits, Fig 2a) and a 1/f background trend (Fig 2a, inset). We then calculated spectrograms of circuit activity to analyze the evolution of signal strength in the time-frequency domain. The spectrograms showed 41 ±3 transient theta-alpha events per 28 s simulation, with average duration of 181 ± 12 ms (Fig 2b). The circuit simulations thus reproduced several key temporal and spectral features of resting-state human EEG, including an oscillatory peak power in theta and alpha bands and a background 1/f trend[49–51]. Critically, these oscillatory properties were emergent, and were not explicitly optimized for, which therefore constitutes an important validation of the model and demonstration of its ability to capture key properties of human cortical microcircuit dynamics.

**Figure 2.**
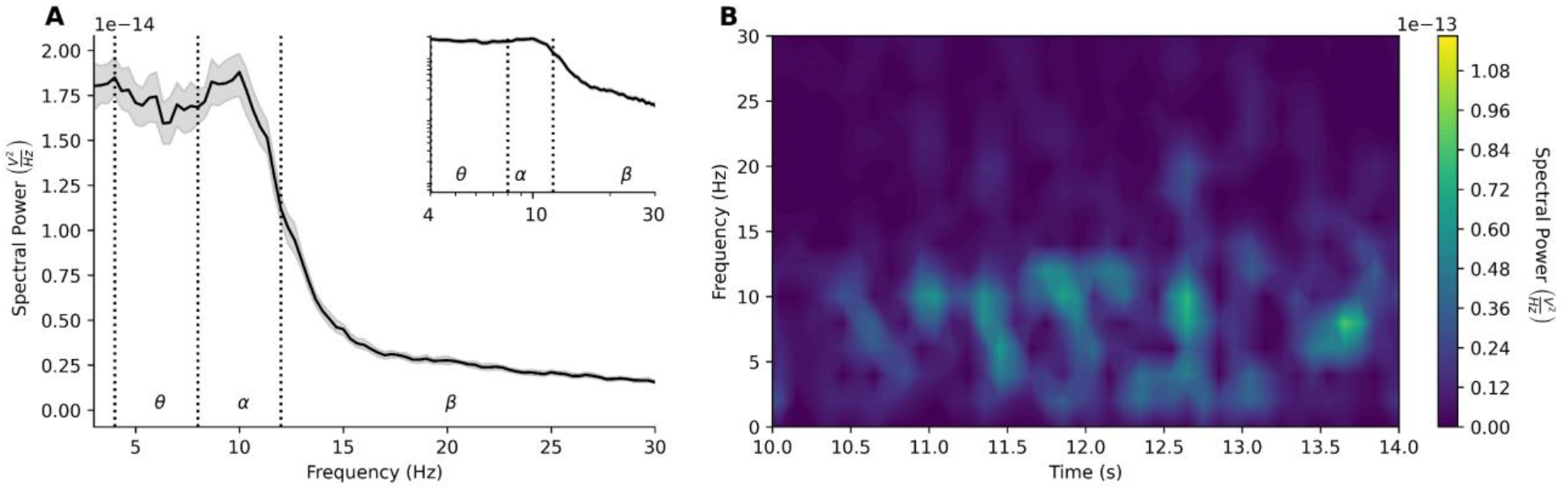
Models of human cortical microcircuits reproduce key features of resting-state EEG. **(a)** Power spectral density plot of circuit simulations (n = 60 randomized microcircuits, bootstrapped mean and 95% confidence intervals), exhibiting peak power in theta and alpha frequency bands. Inset – same power spectral density plot shown on log-log scale, illustrating the 1/f relationship between power and log frequency, inversely linear between 2-30 Hz with slope −0.99 ± 0.07. Frequency bands are delimited by dotted lines. **(b)** Example spectrogram of simulated microcircuit EEG, exhibiting theta and alpha events.

### EEG biomarkers of reduced SST in depression microcircuits

We compared simulated EEG in healthy versus depression microcircuits using our previous depression models, in which SST synaptic and tonic inhibition were reduced (*n* = 60 randomized microcircuits, Fig 3a). The simulated EEG from depression microcircuits exhibited a prominent peak in theta and alpha bands (4 – 12 Hz) similarly to the healthy microcircuits, but there was a significantly increased power in these bands and in low-beta frequencies (5 – 15 Hz, 56% increase on average, *p* < 0.05, Cohen’s *d* = 1.67, Fig 3b). We decomposed EEG PSDs into aperiodic (Fig 3c) and periodic (Fig 3d) components, to compare the distinct functional components of the absolute PSDs. There was a 44% increase in aperiodic broadband power in depression compared to healthy microcircuits (5 – 30 Hz, *p* < 0.05, Cohen’s *d* = 4.9), and a 30% reduction in aperiodic exponent (healthy = 0.99 ± 0.07, depression = 0.69 ± 0.07, *p* < 0.05, Cohen’s *d* = 2.6). The periodic component of the PSDs from depression microcircuits showed a 35% increase in theta power (4 – 8 Hz, *p* < 0.05, Cohen’s *d* = 1.0), and 62% increase in low-beta power (12 – 15 Hz, *p* < 0.05, Cohen’s *d* = 1.9), but no significant change in alpha power compared to healthy microcircuits. Thus, the absolute power increase involved a combination of an aperiodic broadband component, and a periodic theta and low beta component.

**Figure 3.**
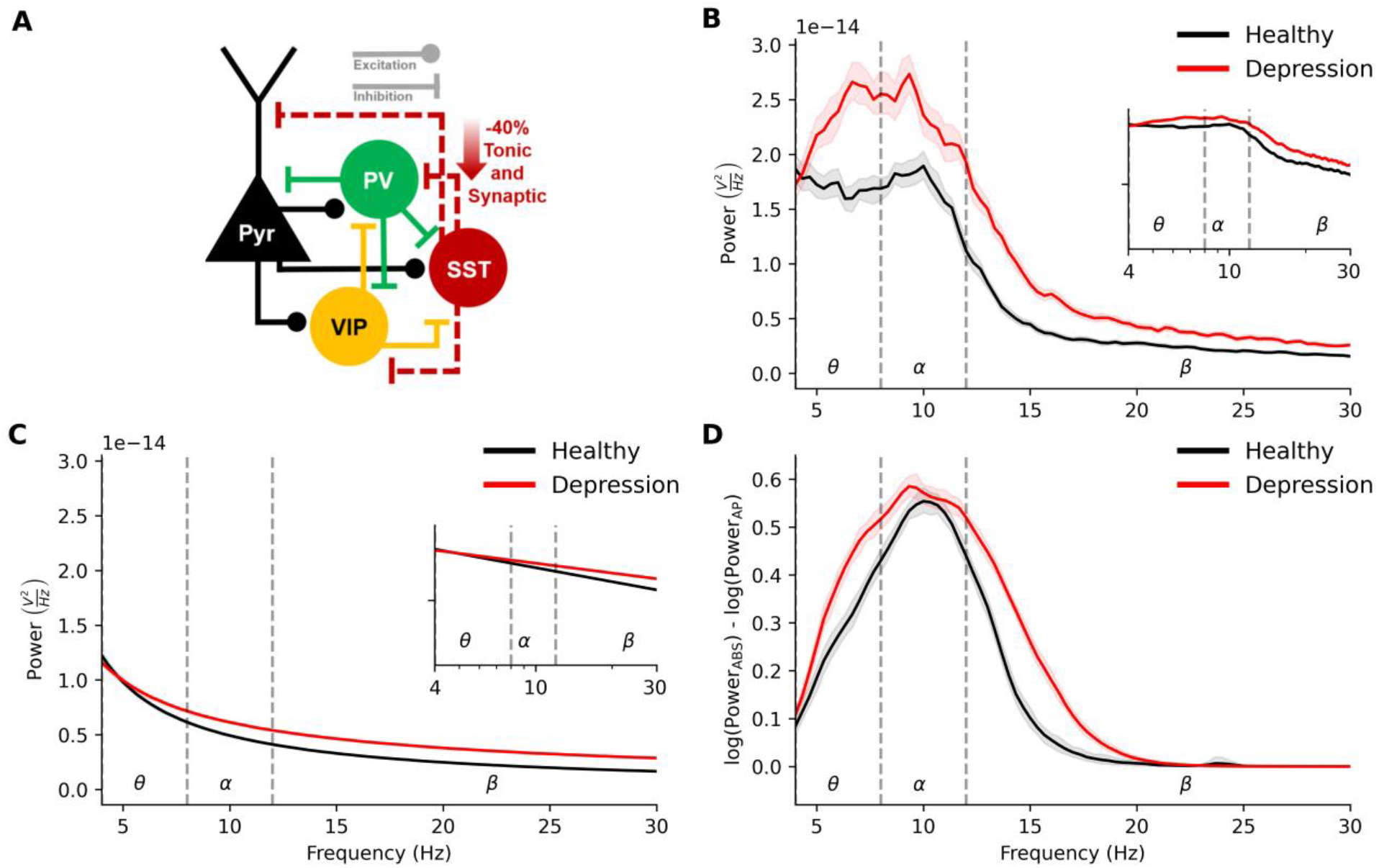
EEG signatures of reduced SST interneuron inhibition in depression microcircuit models. **(a)** Schematic connectivity diagram highlighting the key connections between different neuron types in the microcircuit. In the depression condition, dotted lines from the SST interneurons illustrate reduced synaptic and tonic inhibition onto different neuron types. **(b)** Power spectral density plot of simulated EEG from the healthy (black) and depression (red) microcircuit models (n = 60 randomized microcircuits per condition, bootstrapped mean and 95% confidence interval). (**c**) Fitted aperiodic component of the PSD. **(d)** Fitted periodic component of the PSD.

### EEG biomarkers of reduced SST inhibition are specific

To better establish the specificity of EEG biomarkers due to reduced SST interneuron inhibition, we simulated different levels of SST and PV interneuron inhibition reduction (20-60%) and compared their effects. Spectral profiles of reduced SST interneuron inhibition were distinct from reduced PV interneuron inhibition (Fig 4a). PSDs from microcircuits with reduced PV interneuron inhibition were not significantly different from the healthy microcircuit PSD (20 – 60% reduction, 5 – 15 Hz, 8% change, *p* = 0.3), whereas PSDs from microcircuits with 20 – 60% reduced SST interneuron inhibition had significantly increased power (20% reduction, 5 – 15 Hz, 22% increase, *p* < 0.05, *d* = 0.9: 60% reduction, 92% increase, *p* < 0.05, *d* = 2.3). Following spectral decomposition, we found that the aperiodic component also largely separated the two conditions (Fig 4b). Reduced PV interneuron inhibition microcircuits showed a decrease in broadband power (20% reduction: 8% decrease, *p* < 0.05, *d* = 1.2, 60% reduction: 38% decrease, *p* < 0.05, *d* = 6.2). and no change in aperiodic exponent, whereas reduced SST interneuron inhibition microcircuits showed an increase in broadband power (20% reduction: 21% increase, *p* < 0.05, *d* = 2.9, 50% reduction: 70% increase, *p* < 0.05, *d* = 6.5). In contrast, the periodic increases in power were largely similar between reduced PV and reduced SST interneuron inhibition conditions (Fig 4c, *p* > 0.05 for all reductions).

**Figure 4.**
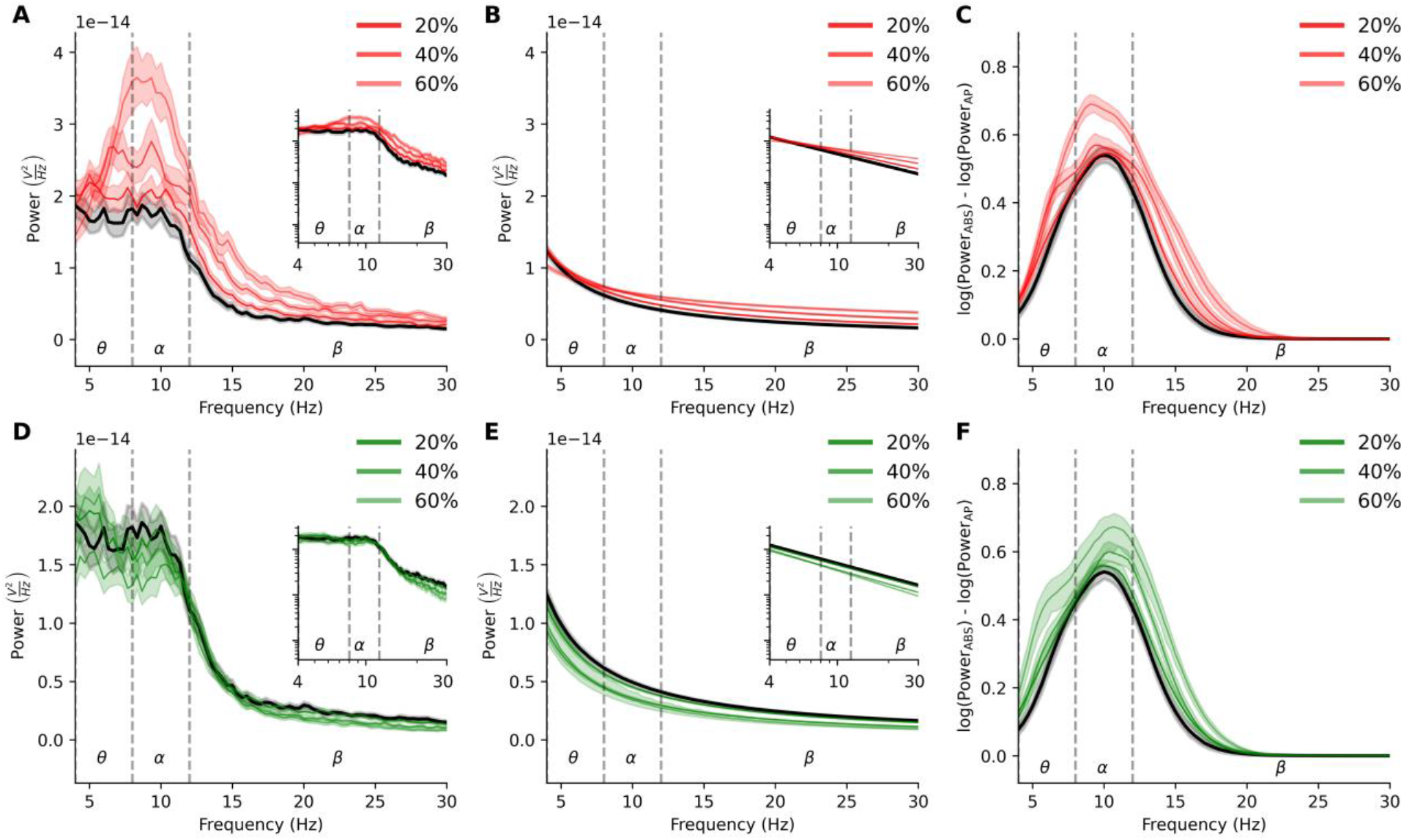
Distinct EEG signatures from reduced SST vs PV interneuron inhibition. **(a)** Power spectral density plot of simulated EEG from the healthy (black) and reduced SST interneuron inhibition (depression, red) microcircuit models, with high-low opacity indicating a 20%, 40% and 60% reduction in inhibition (n = 30 randomized microcircuits per condition, bootstrapped mean). **(b)** Fitted aperiodic component of the PSD. **(c)** Fitted periodic component of the PSD. **(d-f)** Same as (a-c) but for microcircuits with reduced PV interneuron inhibition.

Microcircuits with reduced PV interneuron inhibition significantly increased baseline Pyr firing rate at each step (healthy = 0.84 ± 0.03 Hz, 20% reduction: 0.86 ± 0.04 Hz, *p* < 0.05, *d* = 0.6; 40% reduction: 0.93 ± 0.04 Hz, *p* < 0.05, *d* = 1.7; 60% reduction: 1.12 ± 0.06 Hz, *p* < 0.05, *d* = 4), but to a lesser extent than reduced SST interneuron inhibition (20% SST: 1.03 ± 0.04 Hz, *p* < 0.05, *d* = 5.32; 40% SST: 1.28 ± 0.06 Hz, *p* < 0.05, *d* = 4.8; 60% SST: 1.57 ± 0.08 Hz, *p* < 0.05, *d* = 4.33). Thus, reduced SST interneuron inhibition affected baseline microcircuit activity to a greater extent than reduced PV interneuron inhibition.

We identified relationships between neuronal spiking and EEG spectral changes in depression microcircuit models by determining the EEG phase preference of the neuronal populations. The EEG time series were bandpass filtered from 4 – 12 Hz since the absolute PSDs and the periodic component of the PSDs showed highest power in these frequencies (Fig 5d). Spike time was then converted to phase of EEG signal by temporally aligning the bandpass-filtered signal to the angle of its Hilbert transform (Fig 5a-c). In healthy microcircuits, spiking in all neuronal types exhibited preference to the phase preceding the trough of the EEG waveform (Rayleigh’s *p* < 0.05; Pyr = 269 ± 3°, SST = 267 ± 5°, PV = 283 ± 3°, VIP = 290 ± 2°). Depression microcircuits followed a similar phase preference (Rayleigh’s *p* < 0.05; Pyr = 281 ± 2°, SST = 278 ± 5°, PV = 294 ± 3°, VIP = 296 ± 3°), but there was a decrease in preference selectivity, as quantified by population spike concentrations about the preferred phase (kurtosis, *p* < 0.05 for all; % decrease: Pyr = 10%, *d* = 1.9; SST = 17%, *d* = 1.7; PV = 19%, *d* = 3.2, VIP = 24%, *d* = 4.8).

**Figure 5.**
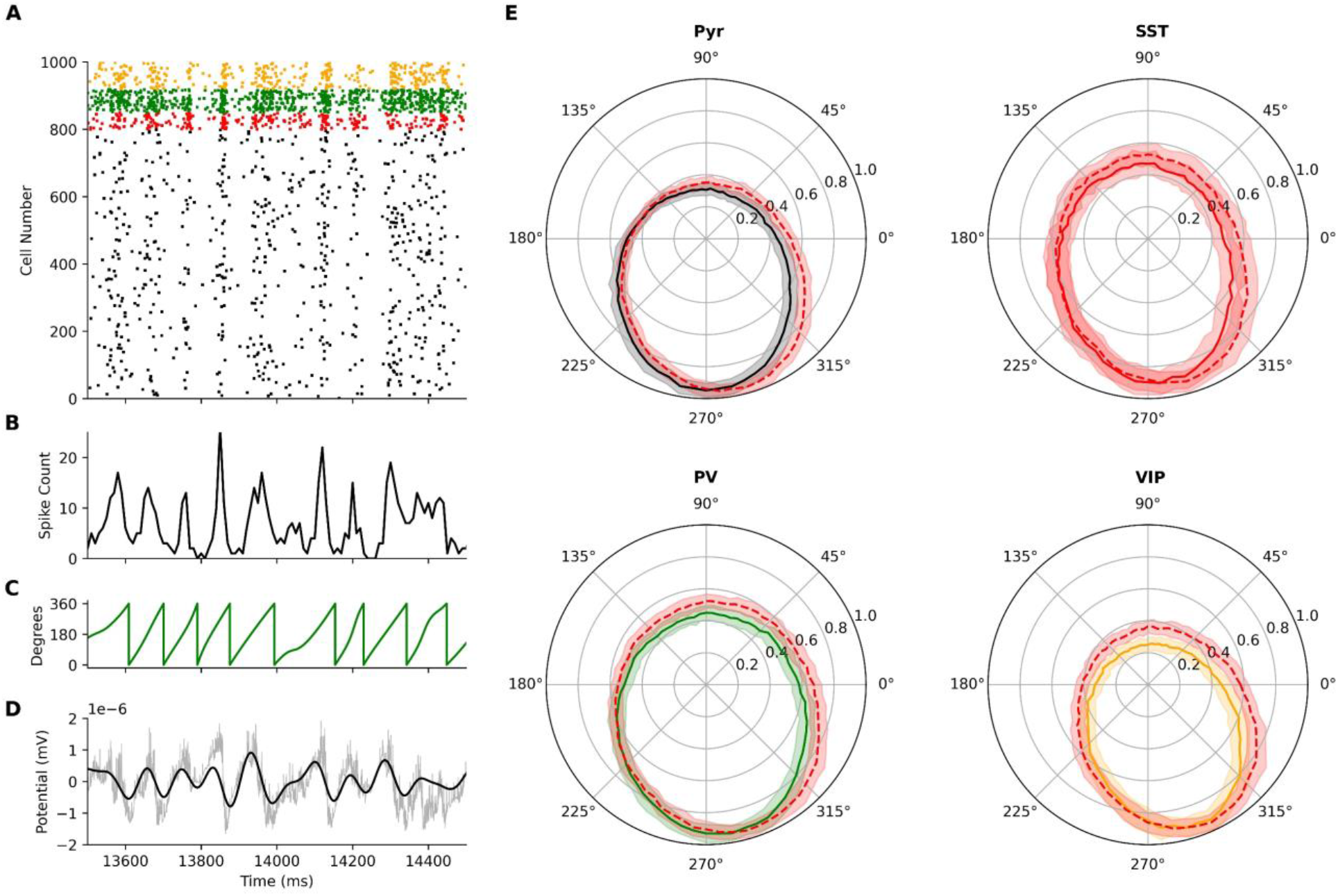
EEG phase preference of neuronal spiking in health and depression microcircuit models. **(a-d)** Temporally aligned simulated microcircuit signals used in calculating population phase-preference. **(a)** Raster plot of spiking in an example healthy microcircuit model. **(b)** Pyr population spike counts, in 10 ms bins. **(c)** Instantaneous phase of the theta-alpha bandpass filtered EEG signal, with trough corresponding to 0°. **(d)** Theta-alpha bandpass filtered EEG signal (black) and unfiltered EEG signal (gray). **(e)** Population spike counts relative to theta-alpha (4 – 12 Hz) bandpass filtered EEG from healthy (solid-line) and depression microcircuits (dotted-line). Plots show bootstrapped mean and standard deviation normalized by peak spike count. Neuron type colors for the healthy microcircuits are as in Fig 1.

Lastly, we examined the spatial distribution of scalp-measured EEG signals generated by the microcircuit simulations using a realistic head model, and determined the distance at which it was indistinguishable from noise. Picking an example region that is also relevant to depression, we placed the simulated microcircuit dipoles into L2/3 of the dorsolateral PFC, at a location directly underneath 10-20 system electrode AF3 (Fig 6a), and used a forward solution based on a boundary-element model with realistic head geometry and conductivity to compute simulated EEG signals. The resulting time series were obtained at different electrode locations on the scalp (Fig 6b). Difference in theta-alpha power in depression versus healthy microcircuits showed a nonuniform decay over the scalp, which was steeper laterally compared to medially (Fig 6c). A significant difference in power was only seen in electrodes corresponding to the ventral portion of the left dlPFC (AF3, the electrode above source) and the left orbitofrontal cortex (FPI and FPz – 29mm and 46mm from source, respectively). The signal spread in both conditions was similarly local and non-uniform, primarily influenced by the location in the head model. Thus, the EEG biomarkers corresponding to microcircuit changes in depression were mostly localized spatially (Fig 6c).

**Figure 6:**
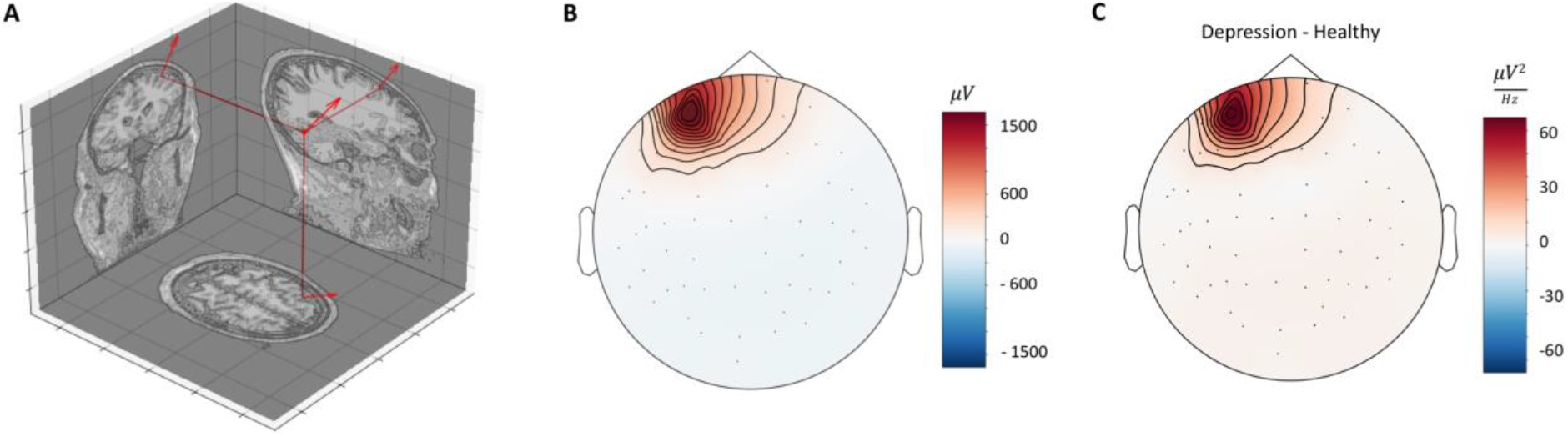
Topographic distribution of microcircuit EEG signal in a realistic head model. **(a)** Microcircuit dipole placement and orientation in the grey matter of the dlPFC in a head model, perpendicular to the cortical surface and in the middle of L2/3 (depth −725 μm). **(b)** Corresponding average potential spread of the microcircuit EEG signal to the EEG electrodes across the head. **(c)** Distribution of the difference in spectral power for depression versus healthy simulated microcircuit signals for theta-alpha band (4 – 12 Hz).

## Discussion

In this study, we identified the signatures of reduced cortical inhibition in depression on EEG signals of resting-state activity using detailed models of human cortical microcircuits. Depression microcircuits with reduced SST interneuron inhibition showed an increase in absolute theta, alpha and low beta power, which involved a broadband increase together with a periodic increase in theta and low-beta activity. These EEG signatures of reduced SST interneuron inhibition were distinct from those corresponding to reducing PV interneuron inhibition, and thus provide a measure of estimating the level of cell-specific inhibition from EEG. These signatures may thus be used as biomarkers to stratify depression subtypes corresponding to altered inhibition, which are particularly associated with treatment-resistant depression[52]. By simulating both spiking and EEG in detailed microcircuits, we showed that neuronal spiking had a marked preference for the phase preceding the trough of theta-alpha EEG band in both conditions. Our results provide mechanistic links between reduced SST interneuron inhibition and candidate biomarkers in EEG signals. These findings have multiple clinical applications, such as improving patient stratification and monitoring the effects of therapeutic compounds targeting SST interneuron inhibition.

Increased power of theta, alpha, and low beta bands in our simulated depression microcircuits is in agreement with several previous depression studies, and was shown to be correlated with diagnosis, severity, and treatment response[7,52–57]. Importantly, systematic reviews across psychiatric disorders showed that these band changes are most consistent in depression and thus particular to this disorder[54]. Further, increased frontal theta power is predictive of second line treatment response targeting cortical inhibition in treatment-resistant depression[52,58–61]. In addition, depression patients who are treatment resistant show especially high theta power compared to treatment responders[52,53,55]. Together with the several points mentioned above, our results provide support for reduced SST interneuron inhibition as a principal mechanism of treatment-resistant depression, and show that it may be detectible by a profile of increased power across the low frequency bands[54,56,62–64]. Future studies relating our simulated EEG biomarkers to experimental EEG and depression features will further establish the link to disease stage, subtype and severity[8,25,65,66]. By incrementally comparing reduced SST inhibition conditions and resulting EEG, we showed that these EEG biomarkers can be used to estimate the level of neuron-specific inhibition noninvasively from patient EEG and thus improve patient stratification. Also, studies may further refine biomarkers by including other mechanisms of reduced inhibition, such SST synapse loss[67]. The increased aperiodic broadband power and decreased aperiodic slope in our depression microcircuits are consistent with the respective effects of increased excitation-inhibition ratio and increased baseline activity[68–70], which in our simulations resulted from reduced SST interneuron inhibition. Increased rhythmic theta and low beta activity, but not alpha, may explain the smaller change in relative alpha power seen previously and lend more functional significance to theta and low beta frequencies[71–74].

Reduced SST and PV circuits both increased baseline activity but to a different degree, and had distinct EEG signatures due to their specific spatial innervation on Pyr neurons[69,75,76] and thus effects on dipole moment generation. The large effect of reduced SST interneuron inhibition on the baseline firing and EEG is supported by the principal role of SST interneurons in modulating baseline cortical activity via facilitating apical dendritic synapses and lateral disynaptic inhibition[25,33,37,77,78]. Changes in SST interneuron inhibition onto apical dendrites of Pyr neuron would therefore have a strong effect on the EEG, since dipole moments increase with distance travelled by axial currents and summate both spatially and temporally[76,79]. In contrast, reduced PV interneuron inhibition had a smaller effect on baseline activity and EEG likely due to the dense PV→PV inhibitory connectivity and non-facilitating synaptic input from Pyr neurons[80], which would dampen the effect of reduced inhibition, and also due to the synaptic innervation of the basal dendrites that would have a smaller effect on dipoles recorded by EEG[76]. EEG decomposition further revealed that altered PV interneuron inhibition had opposite broadband effects on the PSD from SST interneuron inhibition, since decreased inhibitory currents at basal dendrites would negate apical sources and decrease the amplitude of EEG activity[76]. Thus, by decomposing the PSD into its constituents, we further differentiated the absolute EEG signatures of reduced SST from reduced PV interneuron inhibition, highlighting this method’s utility in providing EEG correlates of altered cell-type microcircuitry. These results provide an important validation for our model microcircuits ability to capture a wide range of modulatory effects of distinct inhibitory populations on microcircuit oscillations.

Within physiologically-relevant decreases in inhibition, depression microcircuits increased periodic theta power and decreased theta-phase selectivity. Thus, healthy and depression microcircuits had overall similar oscillatory dynamics, but spiking in the depression microcircuit was somewhat less specific, which is in agreement with leading hypothesis of increased baseline noise in spiking due to reduced inhibition[25]. As previous studies have shown that theta waves modulate network excitability[81], the increased theta activity in depression microcircuits may have further functional consequences via altering cortical dynamics and processing. In summary, our results show that altered cell-specific inhibition in microscale brain circuits in depression is reflected in key features of the EEG spectral density, which can be used to better stratify depression subtype and severity.

Our biophysical models of human cortical microcircuits reproduce the two most ubiquitous features of human resting state EEG: a prominent low-frequency (4 – 12 Hz) peak power and 1/f aperiodic trend[82,83]. In experimental EEG recordings, this low-frequency peak is accentuated when subjects are relaxed with their eyes closed and has the highest test-retest reliability of all spectral features[84]. While recurrent rhythmic activity generated within cortico-thalamic loops is believed to contribute strongly to scalp-measured EEG alpha, cortico-cortical interactions have also been shown to generate alpha activity in top-down processing[85]. Further, it has been shown that cortical Pyr neurons significantly contribute to and sustain theta and alpha-phasic firing due to intrinsic properties and recurrent activity[85–87]. The key features of human EEG reproduced by our microcircuit models were not explicitly constrained for, but rather emerged from the interaction of human neuronal electrical properties, synaptic connectivity, and baseline firing rates. This provides a validation of the models and indicates that they contain circuit motifs responsible for generating realistic microcircuit activity, and it can therefore be viewed as a useful canonical cortical microcircuit model irrespective of layer considerations, in line with previous studies[48]. It will be of interest in future studies to use the models for further elucidating the cellular mechanisms of oscillatory generation, especially in brain states other than rest and during cortical processing, which involve other frequency bands. As our models capture the key spectral EEG features seen *in vivo*, the simulated EEG signal magnitude can also be related to that of the recorded EEG by scaling to account for the difference between the number of neurons in our models compared to the millions of neurons that generate the recorded EEG at a given electrode[36,44].

Linking altered mechanisms at the microcircuit scale to EEG biomarkers is currently impossible to establish experimentally in humans, thereby meriting the use of data-driven detailed computational studies. We simulated human EEG using available detailed models of human cortical L2/3 microcircuits to serve as canonical cortical microcircuit models. These layers are closest to the electrode and thus are important contributors to the EEG signal, along with L5/6[26]. Previous computational studies have shown that interactions between superficial and deep layers are important contributors to electrophysiological signals in other contexts such as event-related potentials[43]. As new human cellular data becomes available, future models that also include deeper layers (4 – 6) could further refine the signatures of reduced SST interneuron inhibition in simulated resting-state EEG signals, by including the dendrites of deep pyramidal neurons and the oscillatory dynamics mediated by interactions between deep and superficial layers[44,88]. Through coupling the biophysical simulations with a forward solution using a realistic head model we showed that EEG signals would be fairly localized to the source and neighboring electrodes. This computational approach can serve to further study source localization in interpreting EEG recordings, improve inverse solutions[89], give greater physiological interpretability to statistical decomposition techniques such as independent component analysis[90] and s/eLORETA[91]. While we modeled and studied a single cortical region at the microcircuit scale, future studies could simulate several distinct microcircuits to examine how altered circuit mechanisms affect multi-regional interactions in depression[25].

Using detailed multi-scale models of human cortical microcircuits, we were able to characterize the EEG and spike correlates of reduced SST interneuron inhibition in depression. Previous modeling studies have shown that reduced SST interneuron inhibition in depression impairs stimulus processing due to increased baseline activity and noise[32]. Our models can serve to identify corresponding biomarkers in task EEG. The computational models we have developed also provide a powerful tool to identify the EEG biomarkers of novel therapeutic compounds and treatments for depression[19,92] via *in silico* simulations to improve treatment monitoring. Finally, our models and methodology may further serve to identify EEG biomarkers of altered cellular and circuit mechanisms in other neurological diseases, such as epilepsy and schizophrenia.

## Methods

### Human cortical microcircuit models

We used our previous models of human cortical L2/3 microcircuits[32], consisting of 1000 neurons distributed in a 500×500×950μm^3^ volume (250 to 1200μm below pia[93]). The model microcircuits included the four key neuron types in cortical L2/3: Pyramidal (Pyr), Somatostatin-expressing (SST), Parvalbumin-expressing (PV), and Vasoactive Intestinal Peptide-expressing (VIP) neurons. The proportions of the neuron types were: 80% Pyr, 5% SST, 7% PV, and 8% VIP in accordance with relative L2/3 neuron densities[94,95] and RNA-seq data[96,97]. The models were simulated using *NEURON*[98] and *ZFPy*[47].

### Resting-state activity simulations

We simulated eyes-closed resting-state by injecting the microcircuit models with background excitatory input representing cortical and thalamic drive, as used previously[32]. The background input was generated by random Orstein-Uhlenbeck (OU) point processes[99] placed on all neurons at the halfway point of each dendritic arbor, and 5 additional OU processes placed along the apical trunk of Pyr neurons in equal intervals, from 10% to 90% apical dendritic length. This enabled the circuit to generate recurrent activity with neuronal firing rates as measured *in vzvo*[31,32,100,101]. For both healthy and depression microcircuit models (see below) we simulated 60 randomized microcircuits, for 28 seconds each.

### Microcircuit models with reduced SST interneuron inhibition (depression microcircuits)

We used previous models of depression microcircuits[32], with 40% reduced synaptic and tonic inhibition from SST interneurons onto all other neurons, in accordance with gene expression studies in SST interneurons in post-mortem brains from depression patients[16]. For Pyr neurons, we decreased apical tonic inhibition by 40%. For each interneuron, we decreased the relative SST contribution to tonic inhibition by 40%. To compare reduced SST from reduced PV interneuron inhibition, we then decreased synaptic and tonic inhibition from SST interneurons by 20%, 30%, 40%, 50%, 60%.

### Microcircuits with reduced PV interneuron inhibition

Reduced PV interneuron inhibition was modeled by iteratively reducing synaptic and tonic inhibition from PV interneurons onto all neurons by 20%, 30%, 40%, 50% and 60%.

### Simulated microcircuit EEG

We simulated layer-averaged dipole moments together with neuronal activity using *LFPy.* Corresponding EEG time series was generated using a four-sphere volume conductor model that assumes homogeneous, isotropic, and linear (frequency independent) conductivity. The radii of the four spheres representing grey matter, cerebrospinal fluid, skull, and scalp were 79 *mm,* 80 *mm,* 85 *mm,* and 90 *mm,* respectively. The conductivity for each sphere was 0.047 S/m, 1.71 S/m, 0.02 S/m, and 0.41 S/m, respectively [102]. A fixed dipole origin was placed at the midpoint of L2/3 (−725 *μm*), and the EEG signal was obtained via forward solution for an EEG electrode placed on the scalp surface. EEG time series were lowpass filtered at 130 Hz. EEG power spectral density (PSD) and spectrograms were calculated using Welch’s method[103], with a 3s second Hanning window for PSD, and 0.5s Hanning window for spectrograms. Both analyses used 80% window overlap.

### EEG periodic and aperiodic components

We decomposed EEG PSDs into periodic and aperiodic (broadband) components using tools from the FOOOF library[68]. The aperiodic component of the PSD was a 1/f function, defined by a vertical offset and exponent parameter. Periodic components representing putative oscillations were derived using up to 4 Gaussians, defined by center frequency (mean), bandwidth (variance), and power (height).

### Population spiking phase preference

We calculated the instantaneous phase of EEG using tools from the *SciPy*[104] and *NumPy*[105] Python libraries. The EEG time series was bandpass filtered from 4 – 12 Hz, with 2 Hz padding on either end to avoid edge effects. A second order Butterworth filter was used to acquire filter coefficients, which were then passed to a linear digital filter applied forward and backwards to eliminate phase lag. The angle of the bandpass-filtered Hilbert transform was calculated and shifted by 180° to obtain the instantaneous phase of the signal with trough and peak corresponding to 0° and 180°, respectively. For each of the four neuron populations (Pyr, SST, PV, VIP), spike times were aggregated into a population spike vector and converted to corresponding phase of EEG. Spike phases were binned into 5° bins and the counts were summated over all circuits in each condition. To compare spike preference between conditions, bin counts were normalized by the maximal count.

### Realistic head source modelling

The x-y-z components of microcircuit dipole moments generated from *LFPy* were imported into *MNE*[106] and downsampled to a 400 Hz sampling rate. The dipole was placed within L2/3 of the left dorsolateral PFC (MNI 30, −36, −42)[107], which has been implicated in depression[108]. We solved the forward solution using a three-layer boundary element model, as implemented in LFPy, corresponding to the inner skull, outer skull, and scalp with conductivities of 0.3 S/m, 0.006 S/m and 0.3 S/m, respectively. The resulting potential was calculated for a standard 60-channel simulated EEG electrode array.

### Statistical tests

We used two-sample t-test to determine statistical significance where appropriate. We calculated Cohen’s *d* to show effect size. We used Raileigh’s test of non-uniformity to determine if populations had a non-uniform phase preference, and kurtosis to quantify variance around the preferred mean angle.

## Data availability statement

All models and code will be available online upon publication.

## Acknowledgements

FM, JG, and EH thank the Krembil Foundation and the Ontario Graduate Scholarship for funding support. FM and EH were also supported by a stipend award from the Department of Physiology at University of Toronto.

## Notes

### Competing Interest Statement

The authors have declared no competing interest.

### Summary of Updates

Revised after peer-review.

